# Comprehensive Analysis of Murine Gait during Skeletal Maturation

**DOI:** 10.64898/2026.02.01.703123

**Authors:** Cameron X Villarreal, Amanda Thayer, Jeremy A. Hampton, Kylie M. Graf, Deva D. Chan

**Author notes:** Corresponding Author: Deva Chan.

## Abstract

Gait is a highly coordinated motor behavior that integrates musculoskeletal and neuromotor control. Analysis of gait parameters is widely used in murine studies as functional biomarkers of locomotor development and disease progression. However, there has yet to be a comprehensive characterization of how gait matures during the rapid growth preceding skeletal maturity, an age range often used in preclinical gait studies. We analyzed gait longitudinally in healthy C57BL/6J mice from 6 to 16 weeks of age, timed to the onsets of sexual and skeletal maturity, respectively. More than 30 gait parameters were quantified weekly and organized into functional groupings reflecting growth, stride, coordination, paw placement, propulsion, and parameter variability. Through a combination of univariate and multivariate analyses, we identified robust age- and sex-based differences across these functional groupings of individual gait parameters. Univariate analysis revealed that many age-associated parameters exhibit a ramp-to-plateau trajectory in the 6- to 16-week age range, with many individual gait parameters plateauing within 8-10 weeks of age. Through multivariate analysis, we identified significant age- and sex-effects on principal components that aligned to functional groupings of gait parameters. Together, these results demonstrate that functional groupings of gait parameters tend to plateau at different phases of skeletal maturation and are differentially impacted by sex, also highlighting the importance of interpreting both univariate and multivariate analysis in longitudinal and comprehensive gait analysis. Understanding these patterns of gait maturation can inform better murine gait study design and more nuanced data analysis that considers potential interactions with age and sex effects.

## INTRODUCTION

Gait is a highly coordinated sequence of motions that enables walking and running, the primary means through which mammals navigate their surroundings, and is disrupted by many diseases, including stroke^1^ musculoskeletal disease and injury,^2,3^ and neurological conditions like Parkinson’s^4^ and Alzheimer’s^5^ disease. Pathological gait disturbances can substantially reduce quality of life,^6,7^ exacerbate disease progression^8^ and are associated with increased fall risk.^9,10^

Because the coordination of complex processes like gait is highly sensitive to pathologic disturbances, evaluating longitudinal changes in gait across maturation provides critical insights into disease progression through gait dysfunction.^11^ For example, Vincelette, *et al*., correlated arthritis severity with stride frequency and length but only characterized gait at 8 and 10 weeks, when growth could obscure pathological effects.^2^ Inoue, *et al*., identified stroke induced alteration in joint kinematics using a single time point in young adult mice.^1^ These studies provide important disease- and development-specific insights, but they do not establish how gait parameters evolve, stabilize, or diverge by sex across the extended period of rapid growth that precedes skeletal maturity. A lack of foundational understanding of gait maturation in healthy mice limits the ability to distinguish pathological gait disturbances from normal developmental variation in gait, even when disease models are compared to age and sex matched controls. Thus, a comprehensive characterization of gait across skeletal maturation in healthy mice is necessary to inform when gait should be measured, how frequently assessments are required and which parameters are most appropriate for detecting disease-associated deviations.

Furthermore, gait varies across age and sex in humans.^12-14^ In mice, however, how a broad range of gait parameters change across age and sex, particularly during post-natal growth and skeletal maturation, remains poorly characterized. Akula *et al*. characterized gait development from 27 to 31 days of age;^15^ however, assessment within a single week limits separation of learning and growth-related effects. This age range is also much younger than those typically used for murine studies of disease progression.^16-19^ Amende, *et al*., evaluated time spent in stance and swing as markers of gait maturation status at 7-8 weeks of age in a Parkinson’s disease model.^20^ Although they evaluated time spent in phases of the gait cycle and several other stance and coordination-associated parameters, how these parameters may be influenced by ongoing growth and skeletal maturation was not considered. Furthermore, skeletal growth is still ongoing at this age, with many bone parameters in mice not considered to be mature until at least ∼3 months old.^21^ As a result, disease-associated gait differences observed during this maturation window may be difficult to interpret without a clear understanding of normal age-dependent gait variation.

Since gait is a complex sequence of spatially and temporally coordinated processes spanning central nervous system control to neuromuscular activation, it manifests as a wide range of measurable parameters. Despite this complexity, studies using gait as a readout often focus on a limited subset of parameters rather than examining gait more holistically. For example, individual parameters describing stride including stride frequency and length are often used to determine if the entire coordinated motion of gait is disturbed.^20,22^ These approaches have yielded important insights, but they rely on a small number of parameters and do not account for the possibility that distinct functional aspects of gait-such as coordination, timing, and execution may mature at different rates or respond differently to experimental perturbations.

Gait changes have been documented across various diseases, but to the best of our knowledge there has yet to be a study that systematically characterizes gait throughout the period of rapid growth preceding skeletal maturation in mice, precisely when gait is studied in many murine studies are studied. This lack of context about gait changes during critical phases of maturation has important implications for experimental design. Without a clear understanding of how various gait parameters change during this window, it is difficult to determine when measurements should be acquired, how to weight sex and age as variables, or which parameters are expected to be stable or highly dynamic during growth. While absolute baseline values will necessarily differ across strains, laboratories, and experimental conditions, the trends in gait maturation and relative stability or variability of specific parameters can provide actionable guidance to study planning. To address this, we performed a comprehensive gait analysis from 6 weeks of age (sexual maturity) to 16 weeks of age (skeletal maturity) in healthy mice applying univariate and multivariate approaches to characterize age- and sex-based dependent changes across individual parameters and functional groupings of parameters. Rather than defining a universal baseline, this study provides a framework to inform experimental decisions that cannot be revisited after data collection, including when gait disturbances are most detectible, which parameters are best suited to capture age- or sex-specific effects, and which parameters are more likely to reflect growth related change versus stable locomotor behavior.

## RESULTS

To characterize how gait parameters change across skeletal maturation, we assessed gait in mice longitudinally from six to sixteen weeks of age timed from the onset of sexual maturity to the onset of skeletal maturity. We permitted mice to run at 20 cm/s on a level treadmill for up to 30 seconds while recording with the Digigait system (Mouse Specifics, Farmingham, MA). Following the manufacturer’s best practices for analysis and similar approaches used by others,^15^ five seconds of video were selected within Digigait analysis software to calculate multiple gait parameters (**Table S1**).

### Functional groups of gait parameters showed distinct dominance of age or sex effects

We grouped gait parameters into seven functional groups (Growth, Coordination, Parameter Variability, Normalized Stride, Absolute Stride, Paw Placement, and Propulsion) based on their contribution to gait (**Table 1**) and tested whether age, sex, or both influenced individual parameters within each group. We found that most functional groups showed a clear dominance of either sex or age effects (**Figure 1**), but Growth and Parameter Variability had equivalent age- and sex-based effects. We determined dominance by the proportion of parameters within each functional group that exhibited a significant association with age or sex (Benjamini-Hochberg-corrected ANOVA q<0.1). Despite the balanced influence of age and sex in the Growth and Parameter Variability functional groups, we detected no significant age-by-sex interaction effects in any individual parameter. Together, these results suggest that age and sex shape gait thorough largely independent, functional group specific means.

**Table 1.**
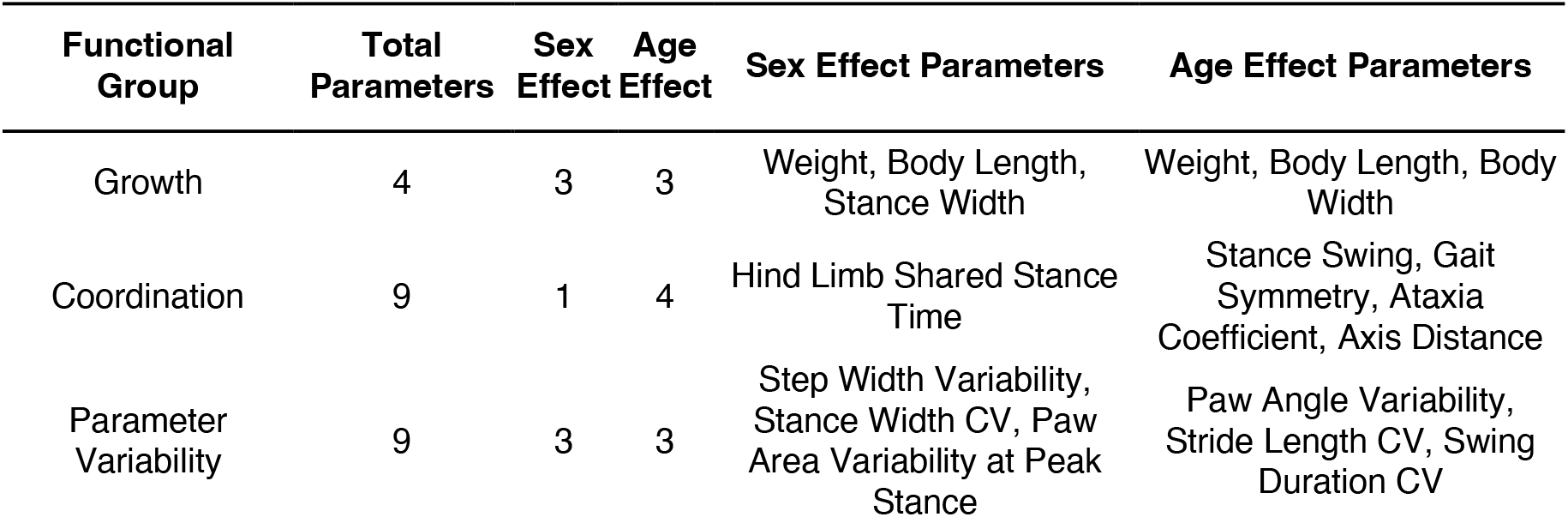
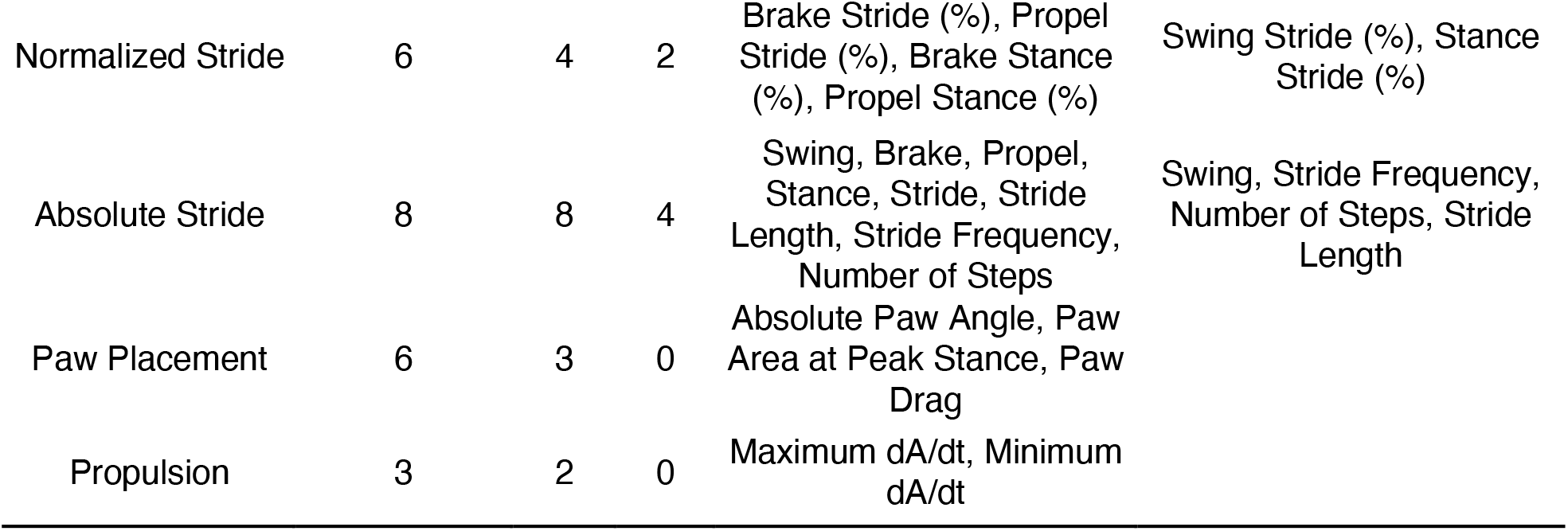
Functional groups of gait parameters that show significant sex or age effects (CV: coefficient of variation; dA/dt: time rate of change of paw area)x.

| Functional Group | Total Parameters | Sex Effect | Age Effect | Sex Effect Parameters | Age Effect Parameters |
| --- | --- | --- | --- | --- | --- |
| Growth | 4 | 3 | 3 | Weight, Body Length, Stance Width | Weight, Body Length, Body Width |
| Coordination | 9 | 1 | 4 | Hind Limb Shared Stance Time | Stance Swing, Gait Symmetry, Ataxia Coefficient, Axis Distance |
| Parameter Variability | 9 | 3 | 3 | Step Width Variability, Stance Width CV, Paw Area Variability at Peak Stance | Paw Angle Variability, Stride Length CV, Swing Duration CV |
| Normalized Stride | 6 | 4 | 2 | Brake Stride (%), Propel Stride (%), Brake Stance (%), Propel Stance (%) | Swing Stride (%), Stance Stride (%) |
| Absolute Stride | 8 | 8 | 4 | Swing, Brake, Propel, Stance, Stride, Stride Length, Stride Frequency, Number of Steps | Swing, Stride Frequency, Number of Steps, Stride Length |
| Paw Placement | 6 | 3 | 0 | Absolute Paw Angle, Paw Area at Peak Stance, Paw Drag |  |
| Propulsion | 3 | 2 | 0 | Maximum dA/dt, Minimum dA/dt |  |

**Figure 1:**
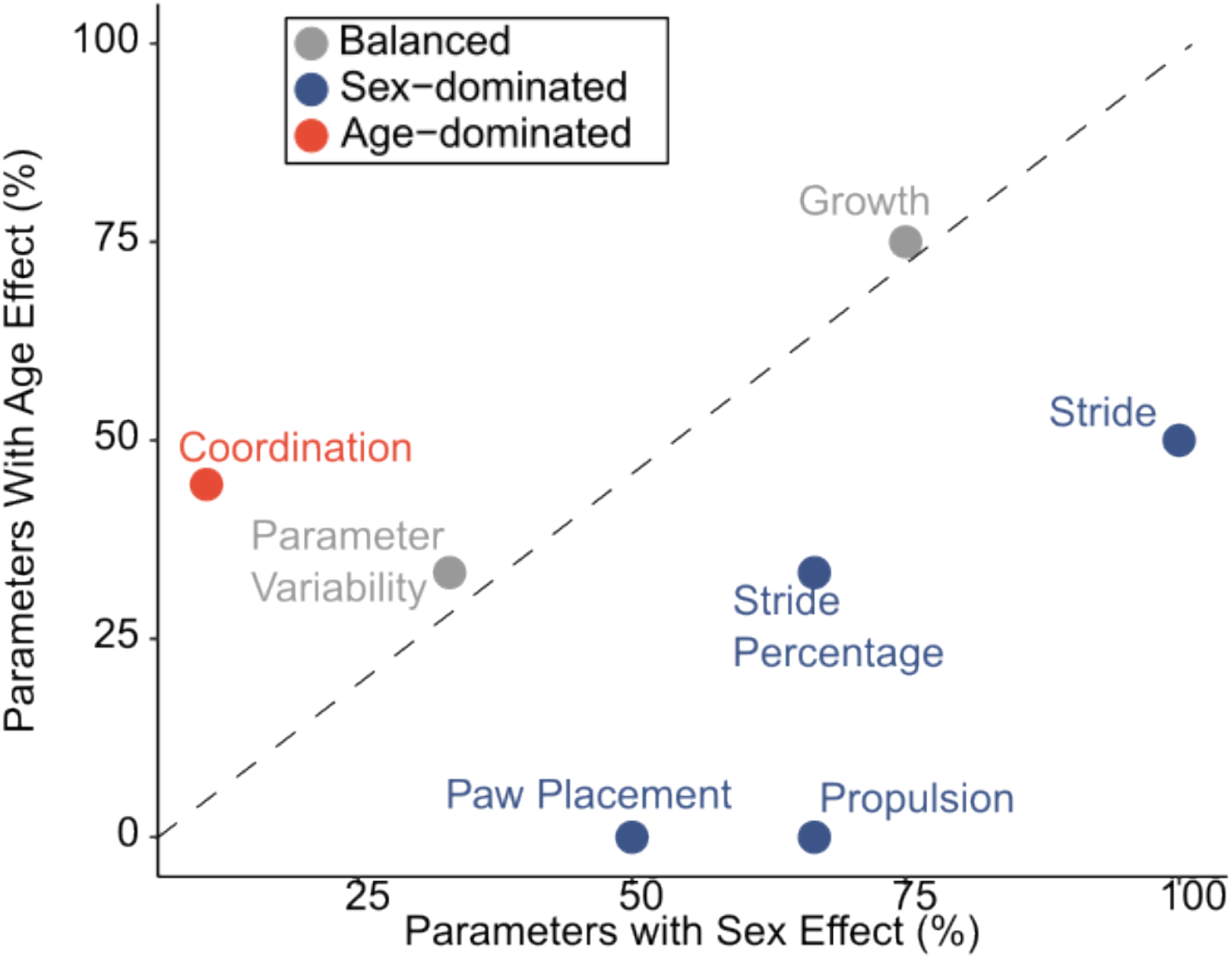
Sex and age differentially shape gait with distinct dominance patterns across functional groups. On this quadrant plot showing the percentage of parameters within each functional group exhibiting significant sex- or age-associated effects, the dashed diagonal indicates equal contributions of sex and age effects. Functional groups positioned to the right of the diagonal are dominated by sex effects, whereas groups to the left are dominated by age effects. Functional groups with a 50 ± 10% split between sex and age effects were classified as balanced. Effect significance was determined using Benjamini–Hochberg–corrected ANOVA, with a false discovery rate threshold of q ≤ 0.1.

### Most age-associated gait parameters plateaued during skeletal maturation

We applied an iterative linear regression approach to parameters with age effects (ANOVA, q<0.1) and assigned a breakpoint week to parameters that showed a clear switch between a ramp phase and a plateau phase. Parameters with other patterns over time were classified as monotonic or sigmoidal, the latter representing a sharper transition between two constant phases. To determine when parameters reached stability, we analyzed percent change from our initial measurement week across week and identified when parameters shifted from active change to fluctuation within the ±95% confidence interval of a stable mean.

Most gait parameters with a significant age effect did not change continuously throughout maturation. Instead, the majority transitioned from an early period of monotonic change (i.e., ramp phase) to a stable plateau that persisted through the onset of skeletal maturity (**Table 2, Figure 2**). Parameters within the same functional group tended to reach stability at similar ages, while different functional groups plateaued at different times (**Table 2, Figure 2**). Parameter Variability metrics stabilized earliest (median plateau: 10 weeks), followed by Coordination parameters (median plateau: 12 weeks), and finally Stride parameters (median plateau: 12 weeks). This sequential pattern suggests that mice first establish repeatable movement patterns, then refine coordination, with both processes stabilizing before reaching adult body proportions. The asynchronous timing of these plateaus indicates that neuromuscular control of gait may mature independently from somatic growth. Sex influenced the timing of gait maturation, but not the magnitude of specific gait parameters. Female mice reached plateau earlier than males for most age-associated parameters (**Figure 2C**). Despite these differences in maturation, both sexes ultimately stabilized with similar mature gait patterns, indicating that the mature gait phenotype converges. These findings suggest that observed sex differences in gait during early maturation likely reflect differences in maturational timing rather than intrinsic sex-based differences in locomotor function.

**Table 2.**
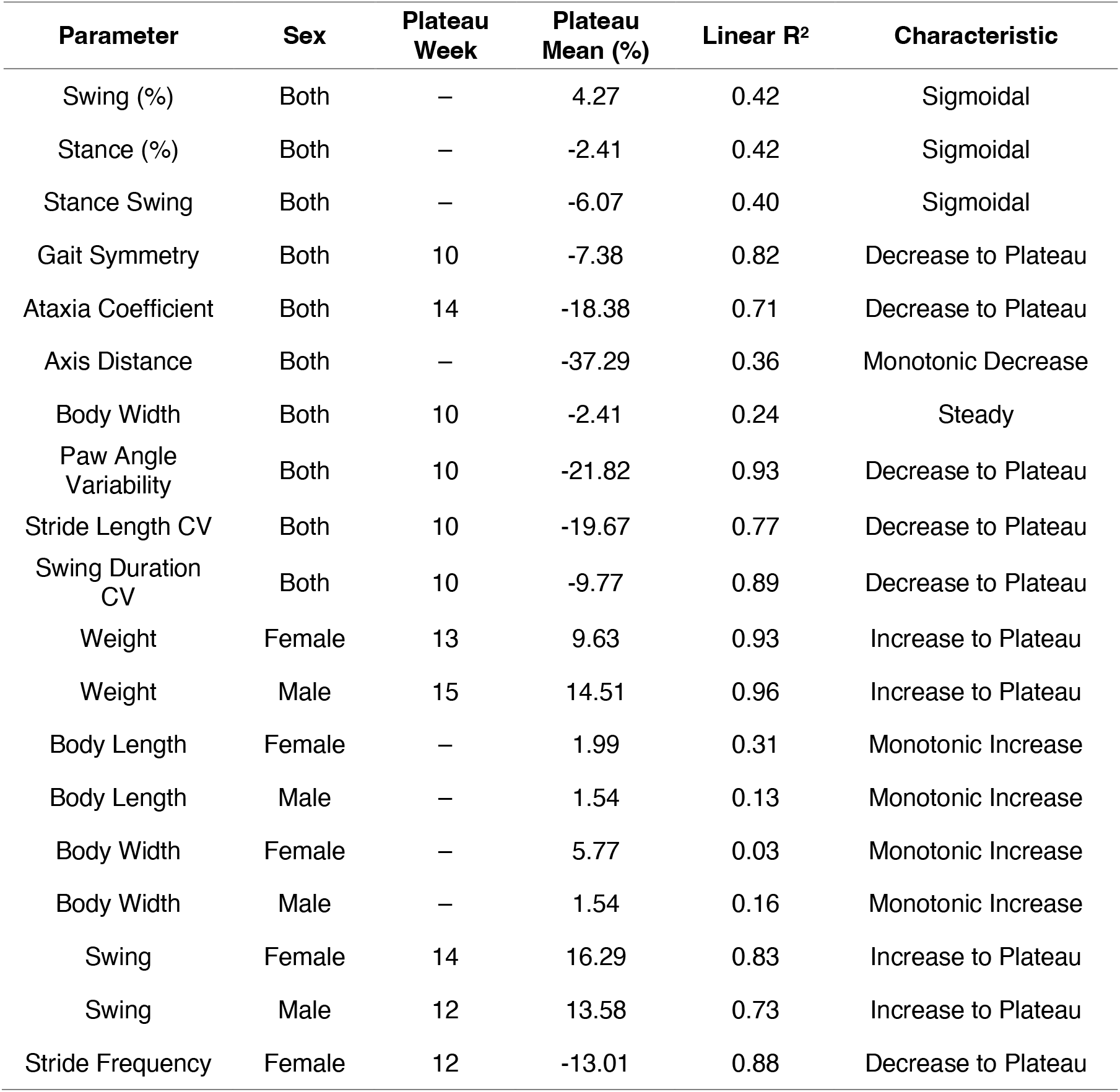
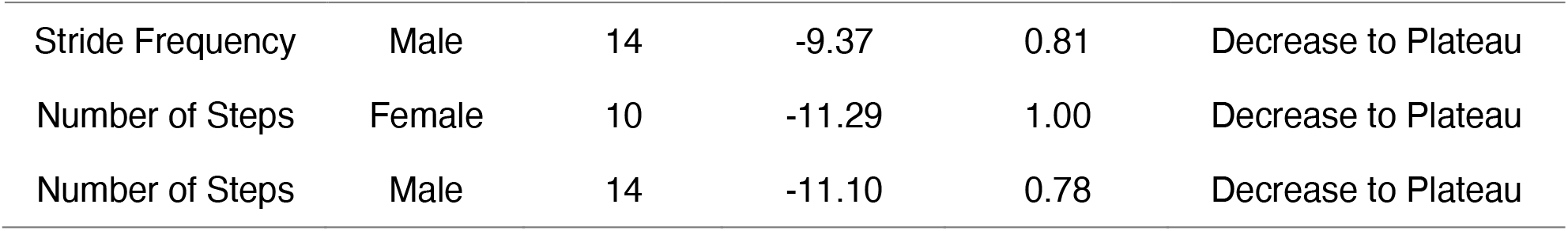
Results of plateau analysis of parameters with a significant age effect. Parameters that have a significant sex-effect detected via Benjamini-Hochberg corrected two way analysis of variance (q<0.1) are separated by sex. Parameters without a significant sex-effect were pooled for analysis. (CV: coefficient of variation)

**Figure 2:**
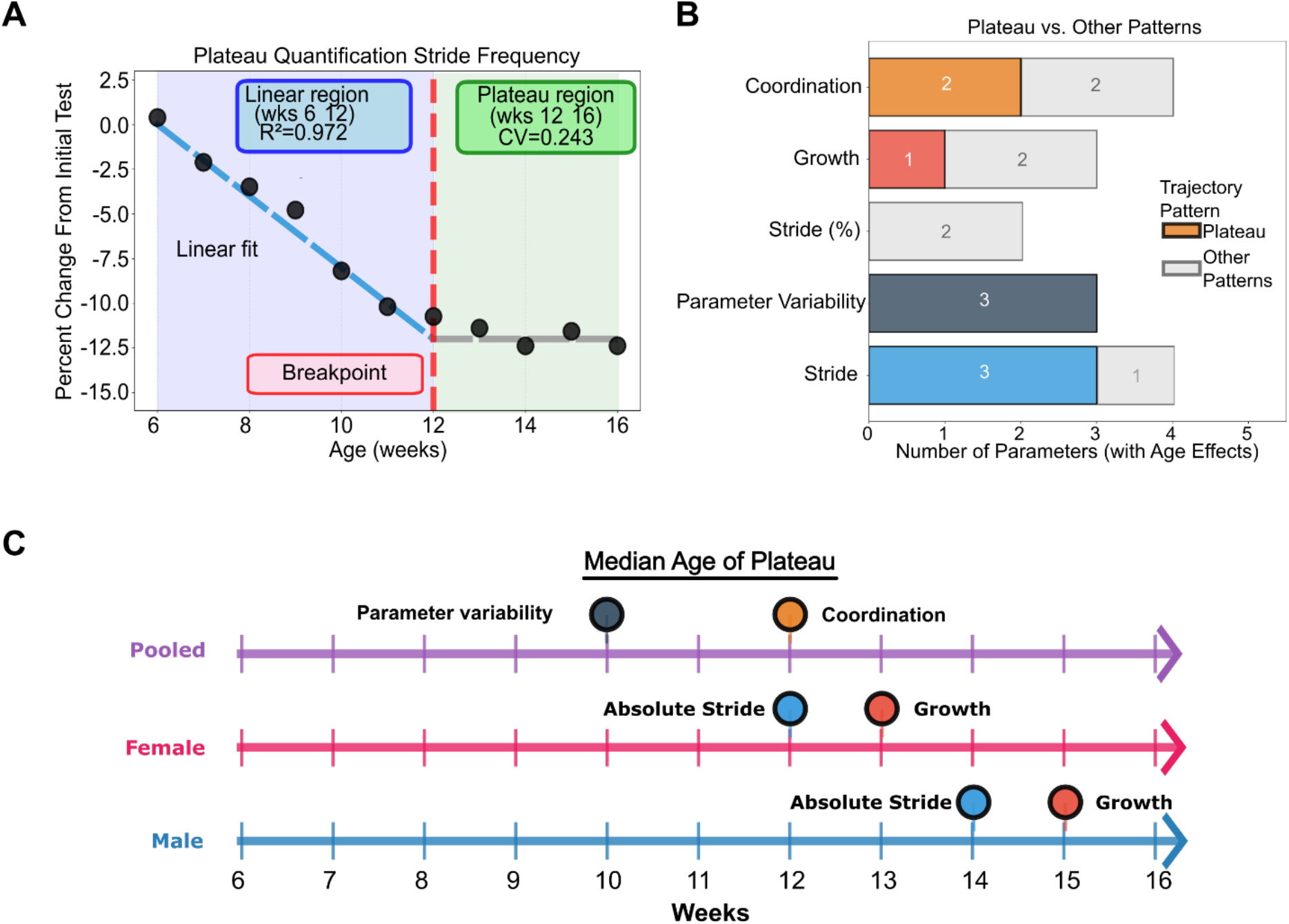
The plateau maturation pattern is the most prevalent growth pattern among individual parameters and this behavior occurs on different timelines for functional groups and across sexes. A) This plot demonstrates the slope to plateau behavior exhibited by parameters with a significant age effect. The blue line represents the linear ramp region, the red dashed line represents the breakpoint, and the grey line denotes the stable mean about which magnitude fluctuates during the plateau period. Values plotted are the percent change in magnitude from the first measurement at week six. B) This box plot shows the number of parameters with significant age effects within a functional group exhibit a plateau analysis. C) This timeline represents the median ages at which functional groups reach plateau, with functional groups labeled. Functional groups with significant sex-based differences are split by sex and functional groups without significant sex-based differences were pooled for analysis.

**Figure 3:**
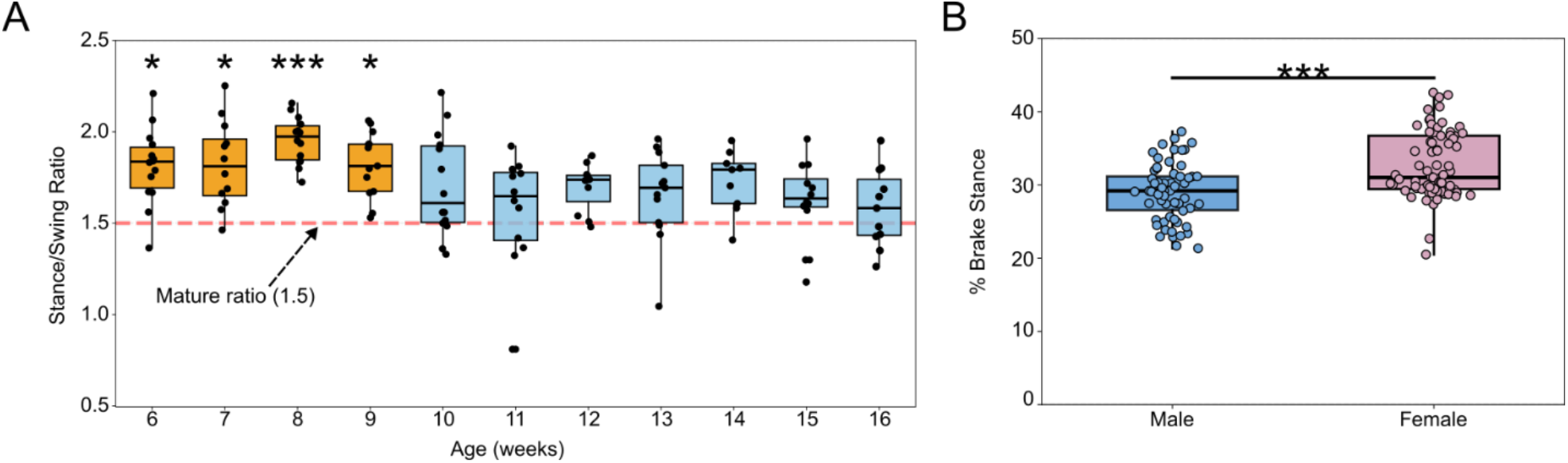
Stance-swing ratio demonstrates maturation effects in gait, while the braking dynamics of stance demonstrate sex-based differences in gait. (A) The ratio of stance to swing for each week was compared to week 16 (skeletal maturity) to identify when this parameter was no longer significantly different from that of a skeletally mature mouse. Mice trended towards a mature stance to swing ratio of 1.5^13,15^ over the course of this study. (B) Males spent a smaller proportion of stance braking than females. The opposite effect was seen in proportion of time spent in propulsion which must sum to 100% with proportion of stance braking. (***, p<0.001).

**Figure 3a:**
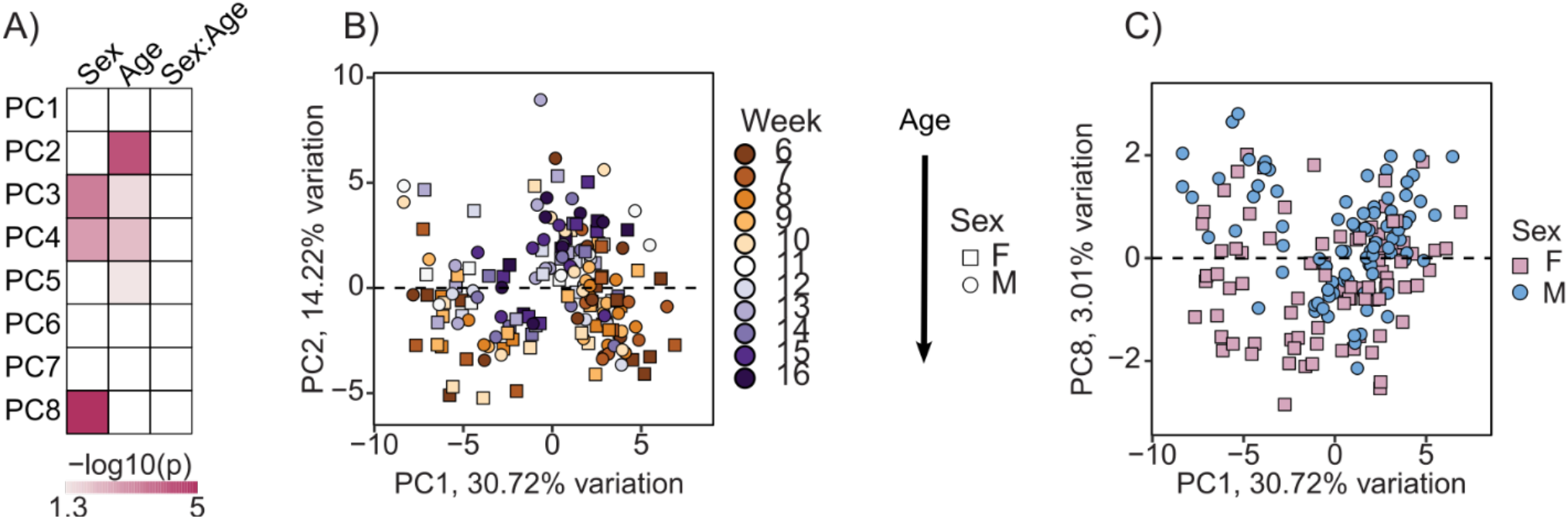
Age- and sex-based effects are linked to specific principal components. (A) A heat map of p values calculated two-way analysis of variance (ANOVA) quantifying sex and age effects on principal components. The intensity of color corresponds to −log_10_(p) with a greater intensity corresponding to significance level. (B) Loadings from individual mice are plotted in PC1-PC2 space, where PC2 is the principal component for which an age effect was most significant (p<0.001). Male mice are indicated with a circle and female with a square; age is denoted along a color gradient from 6 to 16 weeks of age. (C) Scores from individual mice are plotted in PC1-PC8 space, where PC8 is the principal component for which sex was most significant (p<0.001)

### Stance swing ratio stabilized with age, but stance-phase dynamics differed by sex

The stance-swing ratio significantly changed with age while the brake-propel ratio was significantly different between sexes. Because swing (%) and stance (%) exhibited non-linear, but age dependent trajectories, we focused on characterizing that effect. To quantify this maturation pattern, we pooled male and female mice stance-swing ratio at each timepoint. We then compared each time point to week 16, which we used as a reference for mature gait (**Figure 3A**). The stance-to-swing ratio reached a stable value by 9 weeks of age, after which no subsequent time points differed significantly from week 16. This ratio converged toward 1.5, corresponding to a 60/40 stance-to-swing distribution, a hallmark of mature gait reported in both humans and mice.^13,15^

We also identified significant sex-based effects for the percentage of stance spent in breaking and propulsion (Benjamini-Hochberg corrected ANOVA, q <0.1) but these parameters did not show significant age effects. Female mice spent proportionally more time in stance braking, and males spent more time in propulsion (**Figure 3B**). When we pooled data across all time points, a student’s t-test confirmed that both braking and propulsion percentages differed significantly between sexes, indicating that these differences persist across the measured window.

### Principal component analysis revealed distinct age- and sex-associated structure in integrated gait maturation

Having identified age- and sex-dependent effects in individual gait parameters, we next evaluated gait holistically using principal component analysis (PCA) to determine whether these functional group-specific effects persisted when gait was considered as an integrated system. We focused analysis on the first eight principal components (PCs) which together accounted for approximately 80% of the total variance in the dataset (**Figure 3**). PC1 primarily captured broad inter-individual variability across gait parameters, indicated by the absence of an age association and the wide dispersion of PC1 scores across weeks and individuals. The broad distribution of PC1 loadings across multiple functional groups (**Table 3**) indicates that this variability reflects an integrated, whole-gait phenotype rather than differences attributable to any single functional group. We further examined age- and sex-related structure on components with significant effects.

**Table 3:** Relative contribution (%) of functional groups to principal components. Relative contribution was defined as the total loading of each functional group within each individual principal component.

|  | **Coordination** | **Growth** | **Parameter Variability** | **Paw Positioning** | **Propulsion** | **Normalized Stride** | **Absolute Stride** |
| --- | --- | --- | --- | --- | --- | --- | --- |
| **PC1** | 16% | 8% | 18% | 17% | 9% | 5% | 26% |
| **PC2** | 21% | 7% | 14% | 8% | 6% | 26% | 18% |
| **PC3** | 25% | 8% | 21% | 10% | 2% | 23% | 10% |
| **PC4** | 20% | 6% | 22% | 9% | 2% | 34% | 8% |
| **PC5** | 16% | 9% | 28% | 18% | 3% | 19% | 7% |
| **PC6** | 17% | 5% | 28% | 13% | 5% | 6% | 27% |
| **PC7** | 11% | 16% | 32% | 16% | 7% | 8% | 9% |
| **PC8** | 23% | 15% | 14% | 14% | 5% | 3% | 25% |

Among these components, PC2 showed the strongest significant association with age (Pearson’s, r = 0.43, p<0.001) and exhibited the largest age-related effect size (ANOVA, p <0.01). Loadings on PC2 were dominated by parameters from the Normalized Stride and Coordination functional groups (**Table 3**). The most strongly weighted individual parameters (**Figure 4**) included percent time spent in stance (z-scored loading = −1.82), swing (z-scored loading = 2.602), and braking (z-scored loading = −1.54), as well as stance-swing ratio (z-scored loading = −1.89) and shared stance time (z-scored loading = −1.63). These parameters describe gait independently of absolute body size. The structure of PC2 therefore indicates that gait maturation across the measured window is primarily characterized by refinement of stride and interlimb timing rather than scaling effects alone. Other age-associated components, including PC3, PC4, and PC5, showed weaker correlations with age and reflected contributions from Parameter Variability, Paw Placement, and Growth functional groups. Together with PC2, these components demonstrate that age-related variation in gait is distributed across multiple functional groups rather than confined to a single parameter set. Importantly, the functional groups represented within PCs significantly associated with age PCs indicate that gait maturation during the measured window primarily involves coordinated changes in stride time spent in phases of gait and control, with additional contributions from Parameter Variability, Paw Placement, and Growth-related parameters. This pattern highlights the value of evaluating gait using functionally grouped parameters, as maturation manifests through this level of granularity rather than a uniform change across all individual gait features.

**Figure 4:**
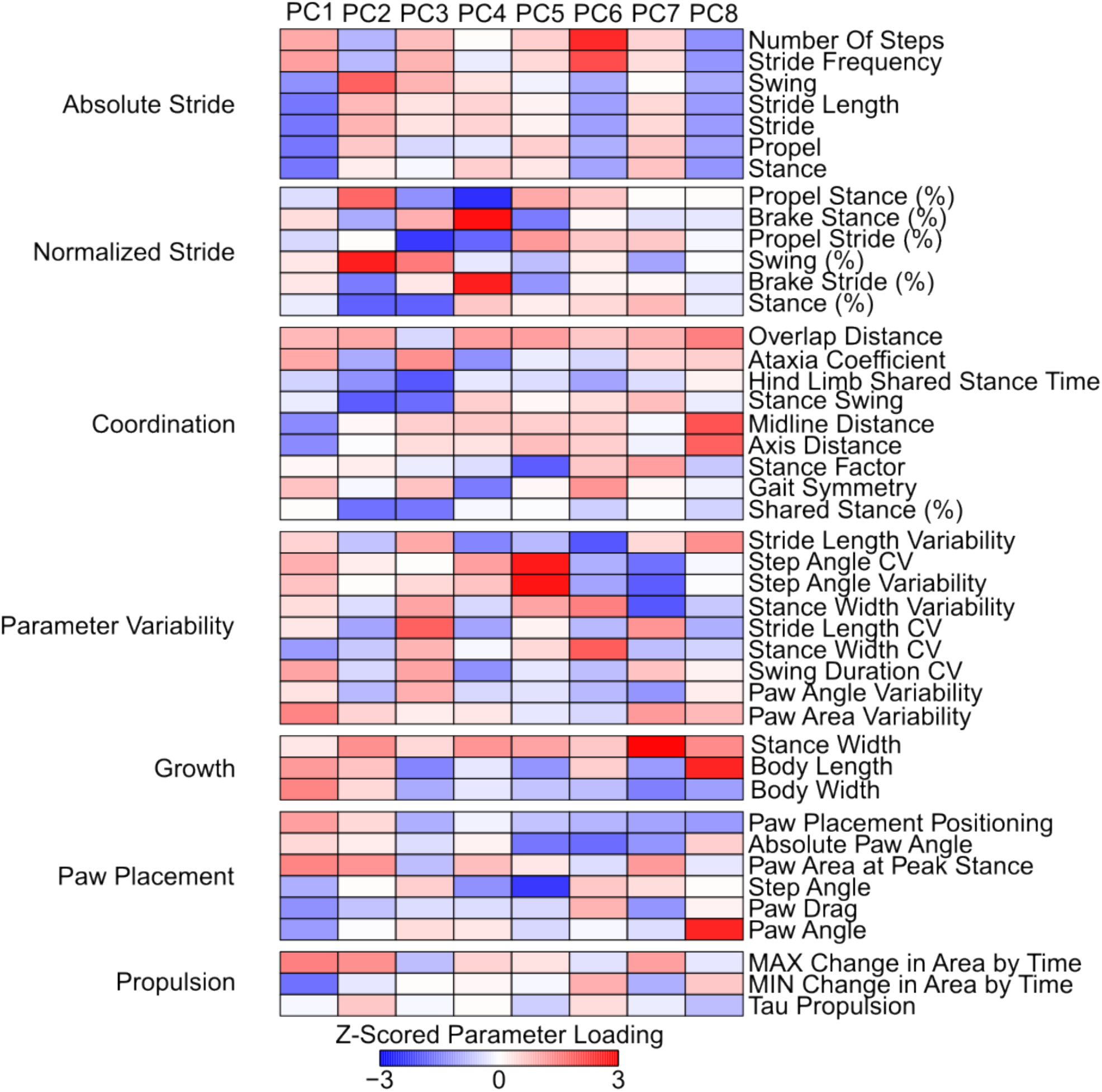
Specific functional groupings of gait parameters drive age- and sex-based differences. A heatmap of z-scored loadings on individual principal components (PCs) demonstrates the magnitude of loading on each functional group. The PCs are arranged in order of decreasing variance explained, while individual parameters are grouped by functional group defined on the left of the heatmap.

We next examined whether sex-based differences observed in individual gait parameters persisted when gait was evaluated as an integrated system. Among the first eight principal components PC8 showed the strongest separation between males and females (student’s t-test. P<0.001). PC8 was dominated by parameters from the Absolute Stride and Growth functional groups (**Table 3**). The most strongly weighted features (**Figure 4**) included time spent in stance (z-scored loading = −1.21) and swing (z-scored loading = −1.00) and body length (z-scored loading = 2.54). These parameters reflect absolute timing and size-related aspects of gait rather than variability or interlimb coordination. Additional components including PC3 and PC4 also showed statistically significant sex differences. Their loading structure reflected contributions primarily from the Parameter Variability and Paw Placement functional groups. Together, these results indicate that combining PCA with functional grouping distinguishes age- and sex-associated gait patterns that may not be apparent from individual parameters alone.

To assess agreement between univariate and multivariate approaches in identifying age- and sex-associated effects across functional groups, we compared the proportion of parameters within each group exhibiting significant age or sex effects with the relative contribution of those groups to PCs showing significant age- or -sex-associations. We identified agreement between univariate and multivariate analysis in only the functional groups for Growth and Absolute Stride (**Table 4**).

**Table 4.**
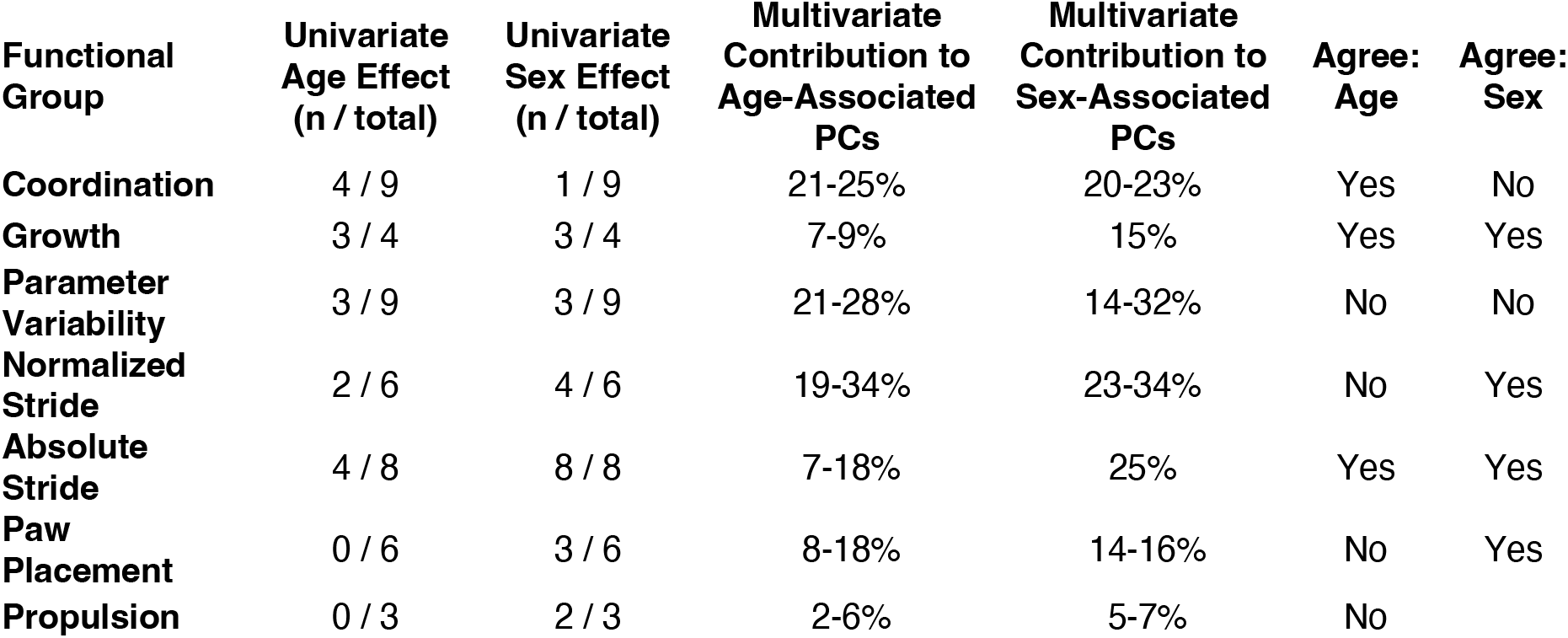
Agreement between univariate and multivariate analyses determined via comparison of significant univariate effects and dominance on sex- or age-associated principal components.

| **Functional Group** | **Univariate Age Effect (n / total)** | **Univariate Sex Effect (n / total)** | **Multivariate Contribution to Age-Associated PCs** | **Multivariate Contribution to Sex-Associated PCs** | **Agree: Age** | **Agree: Sex** |
| --- | --- | --- | --- | --- | --- | --- |
| **Coordination** | 4 / 9 | 1 / 9 | 21-25% | 20-23% | Yes | No |
| **Growth** | 3 / 4 | 3 / 4 | 7-9% | 15% | Yes | Yes |
| **Parameter Variability** | 3 / 9 | 3 / 9 | 21-28% | 14-32% | No | No |
| **Normalized Stride** | 2 / 6 | 4 / 6 | 19-34% | 23-34% | No | Yes |
| **Absolute Stride** | 4 / 8 | 8 / 8 | 7-18% | 25% | Yes | Yes |
| **Paw Placement** | 0 / 6 | 3 / 6 | 8-18% | 14-16% | No | Yes |
| **Propulsion** | 0 / 3 | 2 / 3 | 2-6% | 5-7% | No |  |

## DISCUSSION

In this study, we characterized gait longitudinally in C57BL/6J mice from 6 to 16 weeks of age spanning the onset of sexual maturity through early skeletal maturity. We identified distinct maturation trajectories and sex-based differences that affect both the timing and magnitude of gait features. These results demonstrate that gait maturation in mice is a multi-component process that is better defined by analyzing functional groups of parameters rather than assuming uniform progression across individual parameters. Notably, functional groups of gait parameters plateaued at different time points, and age- and sex-associated effects emerged in largely distinct functional groups. Using complementary univariate and multivariate analyses, we show how individual parameter trajectories and coordinated functional patterns together shape gait maturation. In addition to defining when individual gait parameters stabilize and how age and sex differentially influence distinct functional groups, this work provides practical guidance for the design and interpretation of gait studies in mouse models.

Functional groups of gait parameters plateaued at distinct timepoints (**Table 2, Figure 2**), reflecting asynchronous maturation of different aspects of gait. Among gait parameters with significant age effects, plateau behavior represented the most common maturation pattern across functional groups. The Parameter Variability functional group plateaued at 10 weeks and the Coordination functional group at 12 weeks, with Absolute Stride and Growth functional groups stabilizing later with distinct timings for male and female mice. This maturation sequence suggests that mice first establish repeatable movement patterns, then refine limb coordination, and that both patterns stabilize before achieving adult body proportions. These temporal patterns indicate that gait maturation proceeds through discreet maturational transitions rather than gradual monotonic change.

The predominance of a ramp-to-plateau maturation has important implications for the timing of gait studies. Parameters that exhibit a plateau are less likely to change due to ongoing maturation and therefore provide more stable baselines for detecting pathological effects. In contrast, parameters measured during periods of active change may reflect natural locomotor maturation rather than experimental effect. Accordingly, studies investigating pathologies that affect coordination, such as stroke,^1^ Parkinson’s disease,^4^ or comparing ipsilateral to contralateral limbs in ischemia^23^ and osteoarthritis models,^16^ should strongly consider evaluating baseline gait and initiating experimental conditions after 12 weeks of age to avoid conflating maturational changes and pathological effects. Studies that do not account for locomotor development trajectories risk confounding them with experimental effects.

Individual parameters in the Normalized Stride functional group did not tend to show a plateau pattern. Instead, percent time spent in swing and stance showed a sigmoidal pattern with a trend toward 40% swing and 60% stance (**Figure 2**). In agreement with Akula *et al*., this corresponds to a stance-to-swing ratio of approximately 1.5,^15^ which mirrors the mature gait pattern established in humans.^26,27^ Notably, in humans, this ratio tends to remain constant throughout the lifespan,^26^ but in mice, we observe that this is not the case. Instead, mice in our study reached this stance-swing ratio at 11 weeks of age. This contrasts with Akula *et al*., who identified this split within 4 days of testing,^15^ potentially reflecting both growth and a learning effect. The differing maturation patterns of the stance-swing ratio compared to those in the Absolute Stride functional group suggests an interaction between growth-related scaling and coordination-related refinement that is only evident when gait is evaluated using proportion spent in each phase of gait rather than absolute timing. This distinction indicates that gait maturation cannot fully be captured by absolute timing parameters alone, as percentage phase-based measures reveal coordinated adjustments in how gait is executed that emerge as mice approach skeletal maturity.

Sex-based differences in gait maturation were most apparent in the ramp-to-plateau timing rather than in the ultimate magnitude of specific parameters. Female mice tended to reach stable values than males for several parameters, consistent with their more rapid early growth trajectory (**Figure 2, Table 2**). Despite these differences in maturation timing, both sexes converged on similar mature gait patterns for most parameters, suggesting that early sex differences primarily reflect differences in maturational stage rather than persistent, inherent differences in locomotor control.

Several observed sex effects are consistent with biomechanical consequences of body mass differences,^28^ as male mice are consistently heavier than females across the age range studied. However, body mass alone may not fully account for the observed sex-based differences in braking and propulsion. Human studies have demonstrated that these phases of stance require distinct muscle activation patterns^29^ and regulation of limb loading,^30^ suggesting that sex differences in these parameters may reflect differences in neuromuscular control strategies during gait. Braking and propulsion reflect changes in momentum and are closely linked to mobility^31^ and functional performance.^32^ The persistence of sex-associated differences across the measurement window suggests that these parameters capture sex dependent organization of motor control and muscle activation rather than simple consequences of growth. Accordingly, braking and propulsion provide insight into neuromuscular mechanisms underlying gait control and warrant careful consideration when interpreting gait outcomes in murine models.

While univariate analyses identify parameter-specific age and sex, effects, they may not fully capture how these factors contribute to the complex coordination of gait. We therefore applied multivariate analysis to identify coordinated functional group patterns underlying age- and sex-associated differences in gait. This approach revealed that specific functional groups dominated principal components associated with either age or sex, highlighting coordinated patterns of gait maturation that were not apparent from individual parameters alone (**Table 4**). Multivariate analysis further revealed that some gait features contributed meaningfully to overall gait organization despite showing limited univariate effects. Paw Placement parameters contributed to age-associated components, indicating their relevance emerges through coordinated variation rather than isolated changes to individual parameters. Propulsion parameters contributed consistently to sex-associated components consistent with univariate analysis, reinforcing its role in differential gait execution between male and female mice as opposed to maturation differences. Parameter Variability contributed across most principal components without dominating any single component, suggesting that variability reflects a general feature of gait organization rather than driving age- or sex-specific differences independently. Together these multivariate findings support univariate results by showing that age and sex influence gait through coordinated functional group differences rather than individual gait parameters.

Broader interpretation of these gait characteristics and their sex- or age-linked effects must consider the constraints and limitations of our experimental design. Only a single mouse strain (C57BL/6J) was evaluated; while this strain is widely used, gait maturation trajectories, sex differences, and absolute parameter values may vary between strains. However, we expect the observed trends here to be generalizable given the similar coordination and skeletal maturation timelines across mouse strains. ^33,34^ Gait was assessed using a treadmill-based system with a fixed speed but differs from free ambulation; however, use of a treadmill at consistent settings reduces speed-related variability and facilitates longitudinal analysis. In addition, gait assessments were performed at discrete weekly intervals, which captured broad maturation patterns but may not fully resolve rapid transitional changes occurring between timepoints. However, to the best of our knowledge this is the first study to do this weekly analysis during this time of rapid growth leading up to skeletal maturity. Finally, although analyses were normalized by body width, length, and weight where appropriate, we did not directly assess joint kinematics.^35^ Despite these limitations, the controlled experimental design and consistency between univariate and multivariate findings support the robustness of the maturation patterns we identified during this critical age range for skeletal and gait maturation.

In summary, our findings demonstrate that gait maturation in mice is a multi-faceted process in which age and sex influence distinct functional grouping of gait parameters. We identified distinct plateaus for functional groups of individual gait parameters, where the Coordination and Parameter Variability exhibit a plateau before Absolute Stride and Growth. Absolute Stride and Growth parameters also exhibited distinct male and female patterns, highlighting sex-dependent features that define a mature stride and differentiate gait. These results emphasize the importance of accounting for longitudinal locomotor development and sex when evaluating gait changes, particularly in models of Parkinson’s disease,^36^ stroke,^37^ osteoarthritis,^38^ or other diseases in which degenerative or sex-dependent gait disturbances are expected. Finally, our comprehensive analysis of the week-by-week evolution of gait provides critical baseline information for mice aged 6 to 16 weeks that can guide experimental design of murine gait studies during this critical age range of skeletal maturation.

## METHODS

### Animal Experiments

All *in vivo* murine experiments were conducted under institutional approval (Purdue IACUC protocol 2104002138) and in compliance with ARRIVE 2.0 guidelines. A total of 8 male (from 3 cages) and 9 female (from 3 cages) wild-type C57Bl/6J mice were bred and housed in an AAALAC-accredited facility under a 12-hr light/dark cycle and analyzed at ages between 6 and 16 weeks. Testing occurred during a 2-hour time block in the morning, and mice had *ad libitum* access to standard laboratory chow and sterilized water.

### Gait Analysis

Mice were acclimated to the DigiGait treadmill (Mouse Specifics Inc., Framingham, MA) the day prior to testing to minimize novelty effects. During data collection, animals were placed on the stationary belt, and speed was gradually increased to 20 cm/s, a velocity selected to minimize variability, since speed is a major determinant of gait parameters.^15,39^ That speed was maintained for no more than 180 seconds per mouse across 3 attempts for up to 20 seconds of video capture. Videos were captured from a ventral view using a high-speed camera and analyzed using DigiGait Analysis software (v. 18, Mouse Specifics Inc.), which applies local thresholding and paw-position algorithms to calculate more than 30 stride parameters (**Table S1**) for all four limbs per animal. A usable session was defined as 5 seconds of continuous forward locomotion without slipping or excessive lateral drift. Mice were returned to their home cage immediately following data acquisition. Weekly timepoints without a viable recording were excluded for that mouse. Identified artifacts were corrected according to manufacturer specifications. Stride parameters were averaged across all paws for each mouse at each timepoint for further analysis.

### Plateau Analysis

We applied an iterative linear regression approach to identify maturational plateaus in gait parameters with significant age effects (two-way ANOVA, q < 0.1). For each parameter, we fit linear regressions models to progressively expanding time windows starting from our first measurement at week 6. The initial window spanned weeks 6-8 and was extended by one week per iteration through week 16. At each iteration, we quantified the regression slope and coefficient of determination (R^2^). We defined breakpoints as the earliest timepoint at which the magnitude of percent change shifted from monotonic change to constant with a fluctuation about a mean value within a 95% confidence interval.

### Statistical Analysis

All statistical analyses were conducted in R (v. 4.3.2; R Foundation for Statistical Computing, Vienna, Austria). Gait parameters were first assessed for normality using the Shapiro–Wilk test. We used two-way analysis of variance (ANOVA) to evaluate the effects of maturation or age (6–16 weeks) and sex (female vs. male) on gait parameters, with pairwise comparisons to baseline (week 16) when a statistical effect was detected. A Benjamini-Hochberg correction was applied to ANOVA results to correct for false discovery rate with significance defined as q<0.1. For all *post hoc* pairwise tests, statistical significance was defined as p < 0.05.

We performed principal component analysis (PCA) to reduce dimensionality and identify coordinated patterns of gait development using the PCATools package in R (R Studio version 4.4.2). PCA was conducted on all gait parameters quantified via DigiGait analysis. Prior to PCA, parameters were normalized where appropriate (growth to body length, paw placement to body weight), centered and scaled, and principal components explaining approximately 80% of the total variance were retained for subsequent analyses.

To assess age- and sex-based effects we conducted a two-way analysis of variance on PCs with significance threshold of p<0.05. To assess specific sex-based differences, we used student’s t-test between male and female samples pooled across time points. To assess specific age-based effects we conducted principal components regression with PC score as the response and age as the regressor we evaluated regression coefficients.

To enable fair comparison of gait feature dominance across principal components with unequal variance, we first z-scored parameter loadings within each retained PC and then summarized normalized loadings within functional groups to quantify multivariate dominance, which was compared to univariate age- and sex-associated effects.

## ACKNOWLEDGEMENTS

This work was supported in part by a Defense Advanced Research Projects Agency Young Faculty Award, funded through the Army Research Office as Contract W911NF2110372, to DDC; National Science Foundation Award 2149946 to DDC; National Institutes of Health NIH T32 predoctoral training grant T32DK101001 from the National Institute of Diabetes and Digestive and Kidney Diseases to CXV; and the Indiana Clinical and Translational Science Institute for Indiana University Medical Student Program for Research and Scholarship (IMPRS) support to AT. We acknowledge the assistance of Dr. Wendy Koss (Director of the Purdue Animal Behavior Core) for training and access to the gait analysis system and Ms. Ranya Pendyala for initial work on refining gait analysis protocols. We acknowledge Ms. Aavya Srivastava for initial work refining data analysis pipelines.

## Notes

### Competing Interest Statement

The authors have declared no competing interest.

