## Supplementary material for "Comprehensive Analysis of Murine Gait during Skeletal Maturation": Table S1

**Table S1.** Stride parameters evaluated using the Digigait system. (CV: coefficient of variation; dA/dt: time rate of change of paw area)

| Functional Grouping | Parameter | Unit | Definition |
|---|---|---|---|
| **Absolute Stride Parameters** | Stride Length | Centimeters (cm) | The length that a paw travels in a stride |
| | Stride Frequency | Steps per second | The number of times per second a paw takes a complete stride. |
| | Swing Time | Seconds (s) | The time duration in which the paw has no contact with the belt |
| | Stance Time | Seconds (s) | The duration in which the paw is in contact with the belt. A combination of both propel and brake time. |
| | Stride Time | Seconds (s) | Duration of one complete stride of one paw. The combination of stance time and swing time |
| | Propel | Seconds (s) | The time duration of the propulsion phase when the paw goes from full contact with the treadmill to the start of the swing time |
| | Number of Steps | – | The number of steps for one recording |
| | Brake Time | Seconds (s) | The duration of initial paw contact to full paw contact before the propel phase |
| | Total Number of Strides | – | The total number of strides |
| **Percent Stride Parameters** | Percent Propel Stride | Percent (%) | Percent of the total stride duration where the paw is in the propel phase |
| | Percent Propel Stance | Percent (%) | Percent of the total stance duration where the paw is in the propel stance |
| | Percent Brake Stride | Percent (%) | Percent of the total stride duration where the paw is in braking phase |
| | Percent Brake Stance | Percent (%) | Percent of the total stance duration where the paw is in the braking phase |
| | Percent Swing Stride | Percent (%) | The percent of a total stride in which one paw is not in contact with the belt |
| | Percent Stance Stride | Percent (%) | The percent of the total stride where the paw is in the stance phase |
| **Coordination** | Hind Limb Shared Stance Time | Seconds (s) | Time in which both hind limbs are in contact with the belt |

| | | | |
|---|---|---|---|
| | % Shared Stance | Percent (%) | The percent of total time in which both hind limbs are in contact with the belt |
| | Gait Symmetry | – | The ratio of forelimb stepping frequency to hind limb stepping frequency. |
| | Stance Factor | – | Ratio of left and right stance durations. Right Forelimb to left forelimb and right hindlimb to left hindlimb |
| | Overlap Distance | Centimeters (cm) | Overlap distance between a hind and forelimb pair across successive steps |
| | Ataxia Coefficient | – | (Maximum Stride length - Minimum Stride length)/ Mean Stride Length For one limb |
| | Midline distance | Centimeters (cm) | Distance between the midpoint of the paw (at peak stance) and the transverse midline of the mouse |
| | Stance Swing | – | Stance to Swing Ratio |
| | Axis Distance | Centimeters (cm) | Measure of deviation from the centroid of the mouse during each stride |
| **Parameter Variability** | Paw Angle Variability | Degree (°) | Standard Deviation of the Paw Angle for one set of strides |
| | Paw Area Variability at Peak Stance | Centimeters squared ($cm^2$) | The standard deviation of the Paw Area at Peak Stance |
| | Stride Length Variability | Centimeters (cm) | Standard Deviation of the stride length for one set of strides |
| | Stride Width Variability | Centimeters (cm) | Standard Deviation of the stride width for one set of strides |
| | Step Angle Variability | Degree (°) | Standard Deviation of the step angle for one set of strides |
| | Stride Length CV | Percent (%) | Coefficient of variation of stride length |
| | Stance Width CV | Percent (%) | Coefficient of variation of stance width |
| | Step Angle CV | Percent (%) | Coefficient of variation of step angle |
| | Swing Duration CV | Percent (%) | Coefficient of variation of swing duration |
| | Paw Area CV | Percent (%) | Coefficient of variation of paw area |
| **Paw Positioning** | Paw Area at Peak Stance | Centimeters Squared ($cm^2$) | The area of the paw when the peak stance is measured |

| | | | |
|---|---|---|---|
| | Paw Angle | Degree (°) | The angle that the paw makes with the long axis of the direction of motion. |
| | Absolute Paw Angle | Degree (°) | The maximum paw angle value |
| | Step Angle | Degree (°) | The angle made between left and right hind paws |
| | Paw Placement Positioning | Centimeters (cm) | spatial overlap between ipsilateral fore and hind paw placements during full stance |
| | Paw Drag | | The time-integrated value from full stance to when the paw leaves the belt |
| **Propulsion** | Maximum dA/dt | Centimeters squared per second ($cm^2/s$) | Maximal rate of change of paw area contact during the braking phase |
| | Mininum dA/dt | Centimeters squared per second ($cm^2/s$) | Maximal rate of change of paw area contact during the propulsion phase |
| | Tau Propulsion | Seconds (s) | The time constant of the hind limbs for the exponential decay rate of the gait signal during the propulsion phase of stance |
| **Growth** | Animal Width | Centimeters (cm) | The widest part of the mouse |
| | Animal Length | Centimeters (cm) | The length of the mouse from snout to base of the tail |
| | Stance Width | Centimeters (cm) | The distance between the centers of each axial paw while at peak stance |
